# Integrative transcriptomics reveals sexually dimorphic microRNA control of the cholinergic/neurokine interface in schizophrenia and bipolar disorder

**DOI:** 10.1101/600932

**Authors:** Sebastian Lobentanzer, Geula Hanin, Jochen Klein, Hermona Soreq

**Author notes:** Department of Genetics, University of Cambridge, Cambridge CB2 3EH, United Kingdom (Geula Hanin).

## Abstract

RNA-sequencing analyses are often limited to identifying lowest p-value transcripts, which does not address polygenic phenomena. To overcome this limitation, we developed an integrative approach that combines large scale transcriptomic meta-analysis of patient brain tissues with single-cell sequencing data of CNS neurons, short RNA-sequencing of human male- and female-originated cell lines, and connectomics of transcription factor- and microRNA-interactions with perturbed transcripts. We used this pipeline to analyze cortical transcripts of schizophrenia and bipolar disorder patients. While these pathologies show massive transcriptional parallels, their clinically well-known sexual dimorphisms remain unexplained. Our method explicates the differences between afflicted men and women, and identifies disease-affected pathways of cholinergic transmission and gp130-family neurokine controllers of immune function, interlinked by microRNAs. This approach may open new perspectives for seeking biomarkers and therapeutic targets, also in other transmitter systems and diseases.

## Introduction

Recent large-scale genomic (Anttila *et al*., 2018) and transcriptomic (Gandal *et al*., 2018) analyses uncovered previously unknown magnitudes of transcriptional correlation (71%) in cortical tissues of schizophrenia (SCZ) and bipolar disorder (BD) patients. This suggests a shared SCZ/BD spectrum, but leaves the causes for the widely divergent sex-specific manifestations of these clinical pathologies largely unclear. Compared to women, men present a higher prevalence of SCZ (odds ratio (OR) = 1.4), a ten years earlier mean age of highest disease risk (15-25 vs. 25-35 years of age) and a worse prognosis (Leger and Neill, 2016). In comparison, BD incidence is not dissimilar in men and women, but 80-90% of “Rapid Cyclers” with a particularly bad prognosis are women, and major depressive disorder (MDD), which is a prerequisite for BD diagnosis, affects women more often (OR = 2) (Berger, 2014). Both diseases involve premorbid cognitive impairments and reduced intelligence, but those are more frequent and severe in SCZ than in BD (Bortolato *et al*., 2015), compatible with its original description as “dementia praecox” (premature dementia) early in the 20^th^ century (Kraepelin, 1913).

The genomic origin of SCZ and BD is tremendously complex. Genome-wide association studies (GWAS) have identified several genotype markers distinguishing the (sub-) phenotypes of BD and SCZ (Ruderfer *et al*., 2018) in genes encoding receptors (e.g. for dopamine, glutamate, or acetylcholine (ACh)), scaffolding proteins (e.g. *DISC1*, “disrupted in schizophrenia”), transcription factors (TFs), microRNAs (miRs), or non-coding genomic regions without known function (Harrison, 2015; Henriksen *et al*., 2017; Kanazawa *et al*., 2017). However, the impact of non-coding genomic regions on disease phenotypes was low, possibly indicating that they exert minor effects or do not change expression levels, but rather modify the expression patterns of secondary gene products (Gulyas-Kovacs *et al*., 2018).

Notably, central cholinergic processes are closely associated with disease characteristics and their sexual dimorphism: Male SCZ patients self-administer more nicotine by smoking than females (7.2 vs 3.3 weighted average OR with 90% life time prevalence (Leon and Diaz, 2005)), and pharmacological intervention targeting cholinergic processes is subject to substantial sexual dimorphism (Giacobini and Pepeu, 2018). More specifically, cognitive deficits in SCZ and BD have both been associated with cholinergic dysfunctions (Van Enkhuizen *et al*., 2015; Smucny and Tregellas, 2017) and with the sum of anticholinergic medications (Gray *et al*., 2015; Eum *et al*., 2017), and a polymorphism in the α5 nicotinic ACh receptor associates with both SCZ and smoking in humans, with parallel manifestations in engineered male mice (Koukouli *et al*., 2017) and rats (Forget *et al*., 2018). Correspondingly, cholinergic activation may improve cognition (Sacco *et al*., 2004; Rowe *et al*., 2015; Lewis *et al*., 2017) and mood (Higley and Picciotto, 2014), but provokes schizotypic behavior in Alzheimer’s disease patients (Değirmenci and Keçeci, 2016).

Identifying the cholinergic elements involved in SCZ and BD is challenging. While it is well established that cholinergic projection neurons in the basal forebrain (nuclei Ch1-Ch4) (Li *et al*., 2017) depend on trophic support from their target neurons by retrograde supply of nerve growth factor (NGF) (Levi-Montalcini *et al*., 1996; Mufson *et al*., 2009), not much is known about their cortical counterparts. Other neurotrophic factors repeatedly associated with cholinergic function are the neurokines (McManaman and Crawford, 1991), including Interleukin-6 (IL-6), ciliary neurotrophic factor (CNTF), and leukemia inhibiting factor (LIF). This subgroup of cytokines shares receptors and second messenger pathways. The receptors for IL-6 and CNTF are soluble, secreted proteins, whose signaling depends on binding the other, dimeric transmembrane receptors (Erta *et al*., 2012): the gp130 co-receptor (also known as *IL6ST*, IL-6 signal transducer), and the LIF-receptor. Gp130-family neurokines unite properties of neurotrophic (e.g. on dopaminergic neurons) and immunogenic nature, both prominent facets in the molecular etiology of SCZ and BD (Harrison, 2015) and in cholinergic signaling (Stanke *et al*., 2006). Neurokines can activate *JAK1/2, TYK2* and *STAT1/3/5A/5B* (Rawlings, 2004), affecting neurotransmission as well as immunity. However, to the best of our knowledge, their roles in SCZ/BD were not yet studied.

SCZ/BD hallmarks also involve circadian perturbations. The cholinergic-catecholaminergic imbalance in BD (Van Enkhuizen *et al*., 2015) follows variable transcriptionally-regulated rhythms (e.g. *CLOCK, ARNTL, RORA*), and affected individuals exhibit decreased REM latency (the duration from onset of sleep to the first rapid eye movement phase) and increased vulnerability for disease that can be modulated by muscarinic agonists/antagonists (Ising *et al*., 2005). Correspondingly, the muscarinic *M1/M3* receptor genes are essential for REM sleep (Niwa *et al*., 2018), and sleep deprivation exerts short-term antidepressant effects (Wu and Bunney, 1990), reduced cortical ACh levels (Boonstra *et al*., 2007), and vast transcriptional changes in basal forebrain cholinergic neurons (Nikonova *et al*., 2017).

The quantitative measurement of disease-relevance of individual perturbed genes is a common methodological problem in transcriptomic analyses. To reduce complexity, transcripts are often filtered by their p-values, leading to bias towards few highly expressed and differentially regulated genes. This does not agree with a polygenic model where each component contributes a small effect. To alleviate this bias while still attempting the necessary complexity reduction, we developed an integrative approach of multiple perspectives.

## Results

We first explored the neuronal transcriptomic properties of SCZ and BD in a meta-analysis of deposited cortical male and female patient samples; gene ontology (GO) enrichment analysis of diverging transcripts helped define an objective set of ontological categories to guide further studies. Next, we ascertained co-expression of the corresponding subsystems in cortical tissues by single-cell sequencing analysis, investigating the putative trophic role of neurokine signaling in cortical cholinergic systems and their transcriptional regulation by TFs and miRs. Using pro-cholinergic intervention (stimulation by neurokines) in two closely related cellular models of male and female neuronal origin, we validated the predicted controllers of this molecular interface and identified sexual dimorphisms and affected pathways.

### SCZ/BD transcriptome meta-analysis

#### Sex-independent pathways discriminating between SCZ and BD associate with immunity

Replicating recent reports of sex-independent overlap in patient brain transcriptomes (Gandal *et al*., 2018), we validated the high correlation of SCZ/BD expression beta values (139 SCZ and 82 BD brains with matched controls; Spearman’s rho = 0.7100, p < 0.001). To identify transcript subgroups distinguishing between diseased men and women, we then segregated data of males and females. In both cases, this yielded lower correlations between SCZ and BD than sex-independent data (F: 0.6150, p < 0.001, M: 0.5783, p < 0.001). We sought the most discriminating molecular pathways (i.e., those exhibiting the largest difference in Spearman’s ranks between SCZ and BD), and examined their function. Sex-independently, gene ontology (GO) enrichment of the top 100 diverging genes yielded numerous terms connected to inflammation and immunity (“acute inflammatory response”, p = 0.003, “cellular response to cytokine stimulus”, p = 0.01).

#### Transcriptional sexual dimorphism differs between SCZ and BD

Given the possible male or female biases, we studied all of the 2×2 combinations of the four possible groups (SCZ males, SCZ females, BD males, BD females) by calculating expression beta values inside of each group (against matched controls). We then subjected the most diverging beta values (i.e. the most biased genes) in any meaningful combination to GO enrichment analysis (Figure 1, Data S1). The results indicated a larger divergence between sexes in SCZ than in BD: SCZ-biased genes of males and females showed no overlapping GO terms (Figure 1A), but the top 100 BD-biased genes of males and females showed large GO term overlap, particularly in inflammatory components (Figure 1B). Notably, specific components of neurokine signalling (Rawlings, 2004) were elevated in both males (*IL-6*, p = 0.007) and females (*JAK/STAT*, p = 0.01) with BD.

**Fig. 1.**
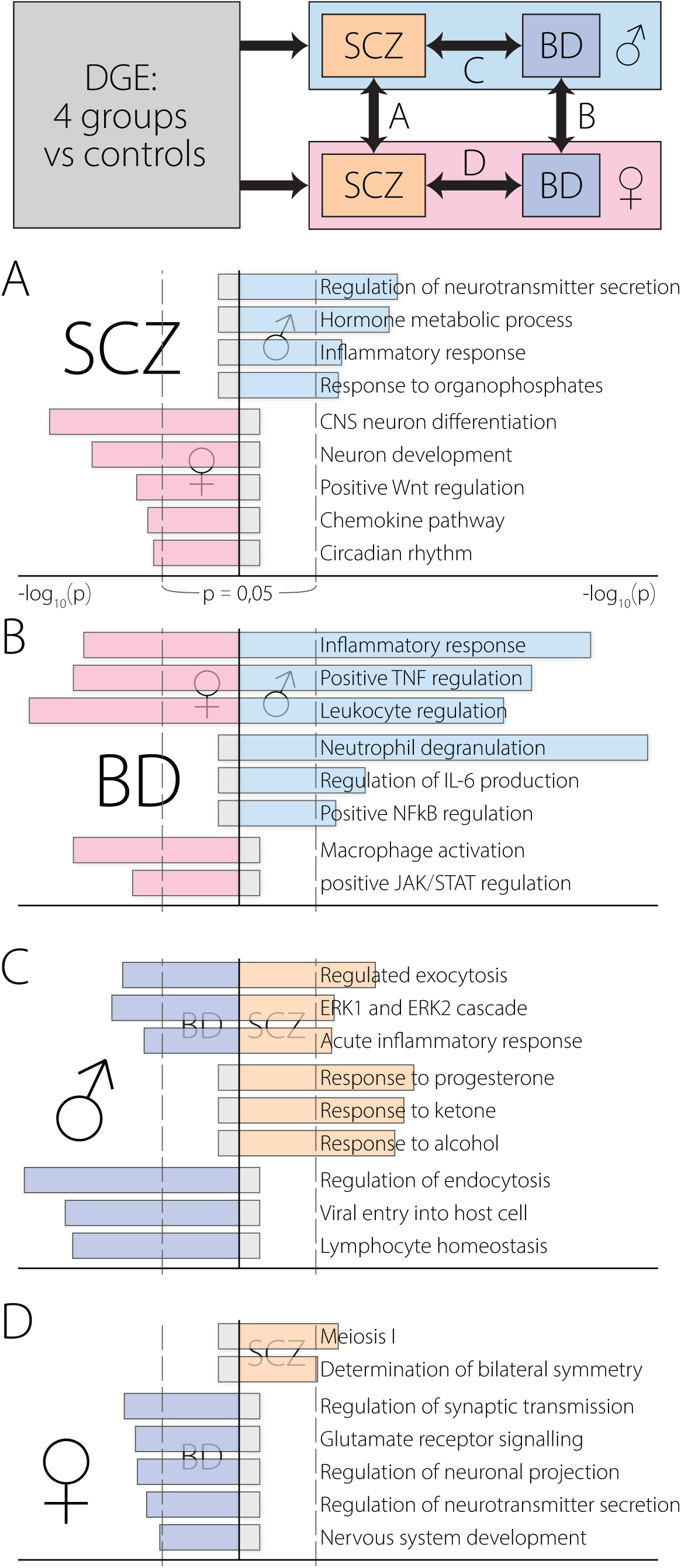
Diverging brain transcriptomes in males and females with SCZ/BD. Differential gene expression results of meta-analysis were dually compared: “SCZ vs BD” or “male vs female”. GO enrichment of the top 100 distinguishing genes in one dimension was compared to the other, for each pair of combinations. (A) SCZ-biased genes diverge between males and females. (B) BD-biased genes share immunological ontology in both males and females. (C) Male-biased genes share immunological ontology in BD and SCZ. (D) Female-biased genes *diverge* between SCZ and BD.

#### Male-biased genes overlap between SCZ and BD

GO terms shared by SCZ and BD patients emerged for male-biased, but not female-biased genes. Males with either disease showed elevated inflammation- and immunity-related genes (Figure 1C). In contrast, female-biased BD genes were enriched in terms associated with CNS function or development (Figure 1D), but enrichment analysis of female-biased SCZ genes yielded no CNS-relevant terms. GO terms pertaining to immune processes were very specific, many referring to single mechanistic components (e.g., *IL-6*), whereas terms focused on neuronal processes failed to implicate specific neurotransmitters. Since the cholinergic systems present significant overlaps with facets of both SCZ and BD, as well as involvement with neurokine signalling (McManaman and Crawford, 1991) and inflammation (Chavan *et al*., 2017), we chose to focus on the diverging cholinergic transcriptomes in males and females with SCZ/BD.

### Cholinergic systems in the CNS

Circos plot analysis of curated functional and anatomical research (Figure 2) and 3D tracing experiments ((Oh *et al*., 2014), Video S1) demonstrate a wide-spread influence of cholinergic systems on both rudimentary and higher cognitive processes throughout the entire mammalian brain (Woolf, 1991; Bina *et al*., 1993; Sarter *et al*., 2009; Mesulam, 2013; Luchicchi *et al*., 2014; Eskow Jaunarajs *et al*., 2015; Gonzales and Smith, 2015; Lin *et al*., 2015; Ballinger *et al*., 2016; Herman *et al*., 2016; Prado *et al*., 2017; Haam and Yakel, 2017; McLaughlin *et al*., 2017). However, trophic factor dependency has only been proven for basal forebrain projection neurons Ch1-Ch4, leaving many open questions about the nature of cholinergic interneurons in the striatum and cortex (Mufson *et al*., 2009) and their transcriptomic features.

**Fig. 2.**
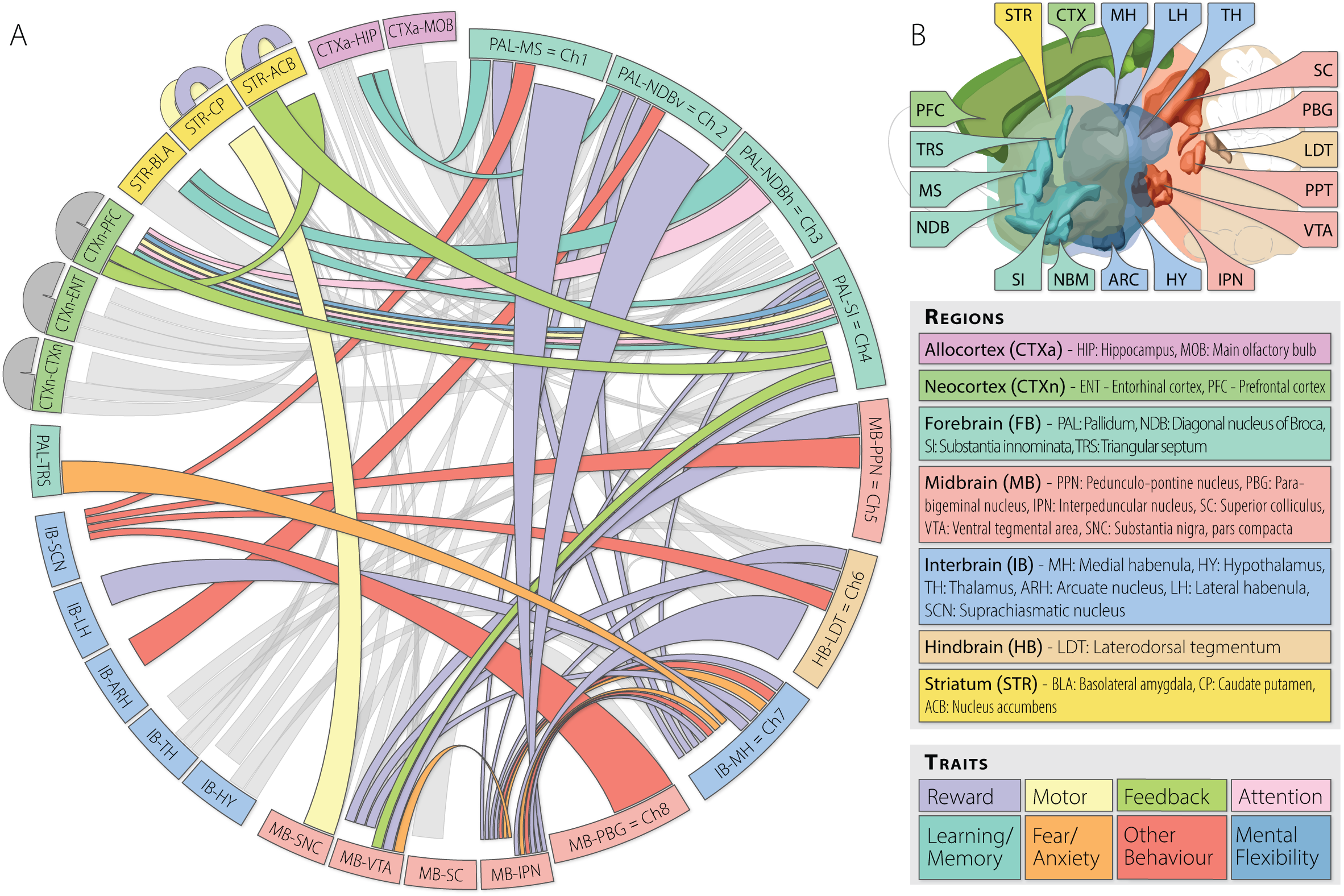
The complexity of cholinergic networks in the mammalian brain. (**A**) Circos overview demonstrating the wide-spread interactions of the mammalian brain’s cholinergic systems with physiologic and cognitive processes. Projection origin denoted by closeness of connector and ideogram (first half clockwise), projection termini by spacing between connector and ideogram (second half clockwise). Cholinergic neurons have been shown in nuclei Ch1-Ch8 (smaller right half of ideogram) and interneurons in the striatum and cortex (outside of ideogram). Functional traits indicated by color of connectors (connector width is determined by geometry and has no implied meaning). (**B**) Brain regions and projection-trait legend for (A). Grey = anatomical cholinergic structure with unknown function.

### Single cell transcriptome analysis

#### Cortical cholinergic cells are mainly neurons and possess neurokine receptors

The predicted transcriptomic interaction between cholinergic and trophic factor systems can only be pathologically relevant if the key elements of both pathways coexist in the same cell. However, patient tissues are almost exclusively collected from brain regions (cortex, hippocampus, seldom striatum) where cholinergic cells are vastly underrepresented (von Engelhardt *et al*., 2007), which complicates the direct retrieval of information on cholinergic processes from total transcriptomes. To examine trophic influences, we turned to web-available single cell sequencing datasets (Darmanis *et al*., 2015; Zeisel *et al*., 2015; Habib *et al*., 2016; Tasic *et al*., 2016) and identified putative cholinergic cells in these datasets as those expressing the cholinergic biomarkers choline acetyltransferase (*CHAT*) and/or the vesicular ACh transporter *SLC18A3* (aka vAChT). Notably, most of these cells or clusters expressed the neuronal marker *RBFOX3* (aka Neu-N), but not the microglial marker *AIF1*. The oligodendrocyte and astrocyte markers *OLIG1* and *GFAP* were both detected in a minority of samples, indicating sparse cholinergic functions in non-neuronal cells. In both mouse and human brains, the identified cells co-expressed the low-affinity neurotrophin receptor Ngfr (aka p75), and the two transmembrane neurokine receptor proteins, gp130 (aka *IL6ST*) and *LIFR* (Figure 3 A-D), but not the high-affinity receptor for NGF, *NTRK1*. In summary, the identified cortical neurons distinguished themselves from basal forebrain cholinergic projection neurons by lacking NGF receptive ability, but possessing the molecular machinery to process neurokine signaling.

**Fig. 3.**
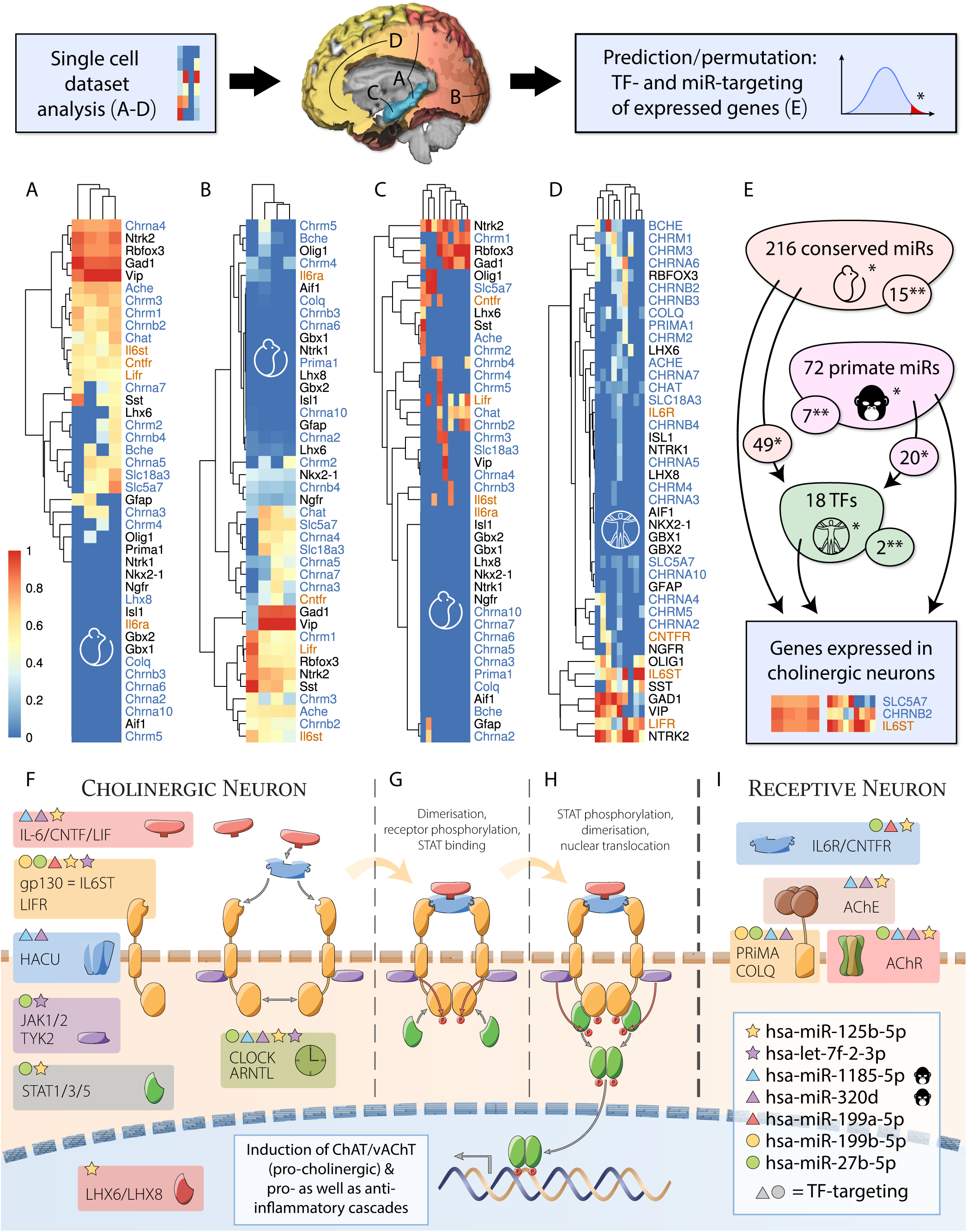
Single cell sequencing of ChAT/vAChT-positive cortical cells and analysis of their expressed transcripts. Expression values were normalized (0-1) for each data set. The order of genes in each heatmap reflects transcript clustering rather than level differences. Columns represent individual samples from original data (column names in Figure S1). (**A)** Clustered single cell sequences of transgenic mouse somatosensory cortex and hippocampus (Zeisel *et al*., 2015); (**B**) Clustered single cell sequences from transgenic mouse visual cortex (Tasic *et al*., 2016); (**C**) Single nucleus sequencing of adult mouse hippocampus (Habib *et al*., 2016); (**D**) Single cell sequencing of human developing neocortex (Darmanis *et al*., 2015). Cholinergic genes denoted in blue, neurokine receptors in orange. (**E**) Permutation analysis of miR- and TF-targeting data of genes expressed in cholinergic cells (via *miRNet*) identified putative cholinergic/neurokine co-regulators (*: p < 0.05, **: p < 0.001). Implicated TFs are regulated by a subset of miRs targeting cholinergic genes, indicating nested regulation (details in Data S2). **(F-I)** Gp130-family neurokine, cholinergic, and circadian signaling pathways are controlled by primate-specific and evolutionarily conserved miRs. miR-targeting of individual genes (colored nodes) yields complex transcriptional interaction. Several miRs directly targeting the cholinergic pathway also target TFs controlling this pathway (circle and triangle).

#### Neuronal transcripts predict regulatory transcription factor and microRNA circuits

TFs exhibit particularly high complexity in the central nervous system (Marbach *et al*., 2016), as do short and long non-coding RNAs (ncRNAs), regulatory elements whose expression is modified in mental diseases (Harrison, 2015). The best studied ncRNA species are microRNAs (miRs), small (18-22 bases) single-stranded RNAs which interfere with translation of transcribed mRNA via guiding an RNA-Induced Silencing Complex (RISC) to the mRNA through sequence complementarity followed by inhibition and/or degradation of those mRNAs. By targeting multiple genes, miRs can exert complex contextual regulation in a temporal and/or spatial fashion (Greenberg and Soreq, 2014), retaining homeostasis, which is paramount for nervous system health (Shaltiel *et al*., 2013), or leading to nervous system disease when disturbed (Bekenstein *et al*., 2017; Rajman and Schratt, 2017).

To pursue the coding and non-coding regulatory elements of cholinergic/neurokine processes, we employed targeting analyses of TF-gene and miR-gene interactions in a graph database specifically constructed from comprehensive state-of-the-art data (*miRNet*, see Methods), using random permutation to empirically estimate false discovery ratio (FDR). This identified 288 miR candidates (out of all 2588 annotated mature miRs) and 18 TFs (out of 618) with FDR < 5%, and 22 miRs and 2 TFs with FDR < 0.1%. The indicated TFs are controlled by a subset of the identified miRs (69 predicted, 12 of those with experimental support), indicating nested layers of regulation (Figure 3E, Data S2). This gave rise to the hypothesis outlined in Figure 3F-I, whereby a cooperative pathway regulation by miRs and TFs connects cholinergic neuronal function with endocrine or paracrine trophic signalling through neurokines. Together, these analyses called for experimental validation of the indicated regulatory pathways linking TFs, neurokine-cholinergic signaling and miRs, in male and female cells.

### Short RNA profiling

#### Neurokine-induced cholinergic differentiation distinctly alters short RNAs in male and female human neuronal cells

Identifying sexually dimorphic miRs of cholinergic relevance in homogenized patient brain samples is challenging due to their tissue-specific and low-level expression (Liu *et al*., 2016). As an alternative, we used immortalized cell lines of male and female human neuronal origin which undergo cholinergic differentiation when subjected to neurokine stimulation (McManaman and Crawford, 1991). Briefly, we exposed the female cell line LA-N-2 and the male cell line LA-N-5 to CNTF, and used short RNA-sequencing (GSE132951, 4 biological replicates) to identify the affected miRs following exposure to this neurokine (Figure 4A). Both cell lines exhibited an immediate response of miR expression as early as 30 minutes after CNTF onset, with increasing numbers of differentially expressed (DE) miRs under prolonged CNTF exposure (Figure 4B, Data S3). In total, we detected 490 DE mature miRs: 107 in LA-N-2, 269 in LA-N-5, and 114 in both. Notably, the female-originated LA-N-5 cells responded more strongly to the neurokine stimulus, showing more DE miRs with a trend towards higher count-change values (mean of absolute count-change across all DE time points, 20907 vs 3066, p = 0.08).

**Fig. 4.**
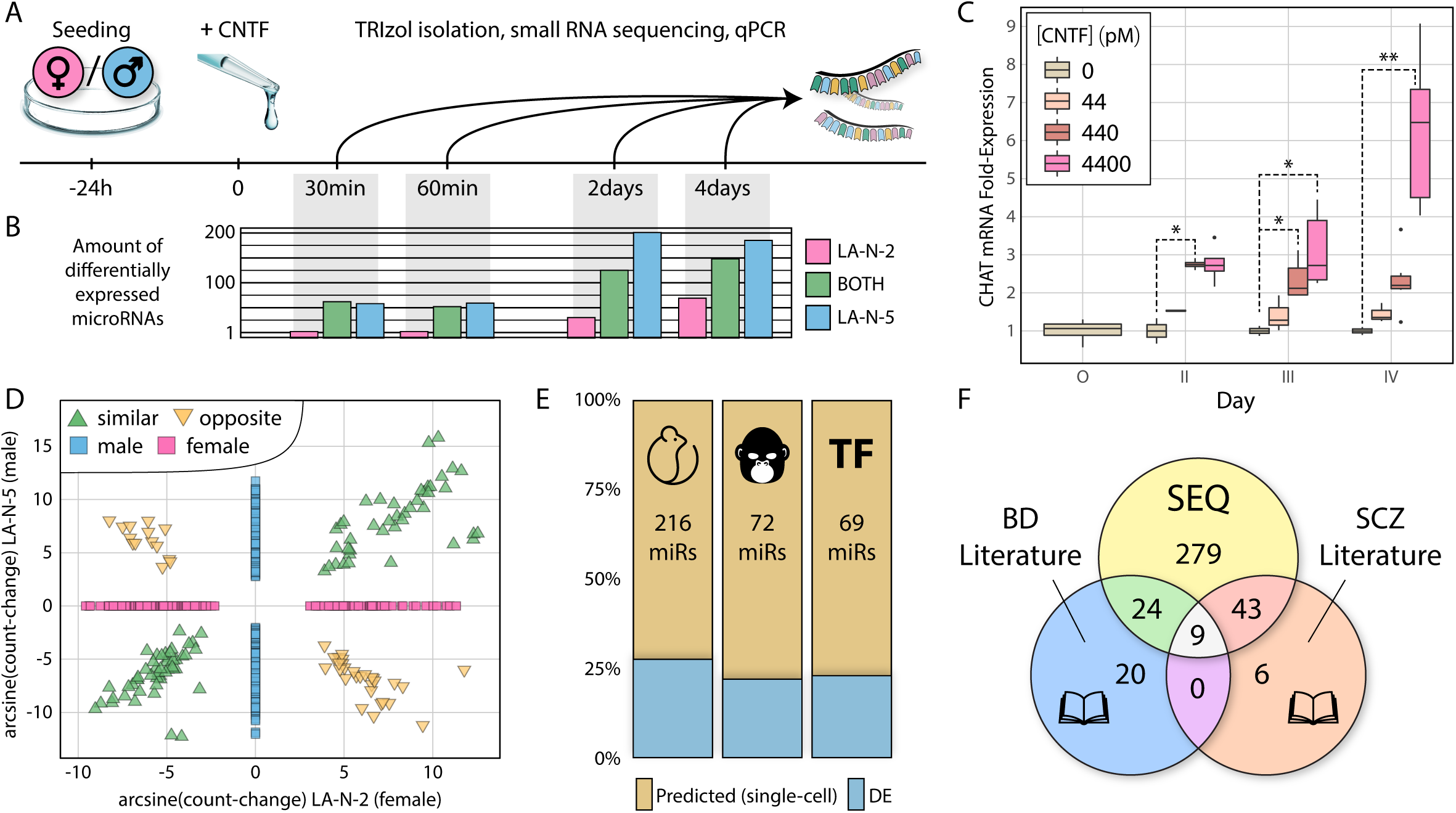
Cholinergic response to CNTF induces partially overlapping miR changes in female- and male-originated LA-N-2 and LA-N-5 cells. (**A**) CNTF differentiation of cells, timeline. (**B**) Bar graph of miRs differentially expressed (DE) during exposure of LA-N-2 cells to 4,4 nM CNTF (pink), LA-N-5 cells to 440 pM CNTF (blue), or in both (green). (**C**)Time-dose boxplot of *CHAT* mRNA expression in LA-N-2 during differentiation with CNTF. For LA-N-5, see Figure S2. *: p < 0.05, **: p < 0.001. (**D**) DE miRs overlap between LA-N-2 and LA-N-5. miRs DE same- and opposite-directionally show highly correlated count-changes (arcsine transformation used because negative values prohibit log-fold display). (**E**) Neurokine-induced DE miRs partly overlap conserved, primate-specific, and TF-targeting miRs predicted via single cell analysis (Figure 3E). (**F**) DE miR precursors overlap with known BD- and SCZ-relevant miR precursors (Beveridge and Cairns, 2012; Fries *et al*., 2018).

Pico- to nanomolar concentrations of CNTF, well within its physiological range (Sun *et al*., 2016), elevated the cholinergic marker genes *CHAT* (P_Day3_ = 0.005, P_Day4_ = 4.1E-05) and *SLC18A3* (aka vAChT, P_Day2_ = 0.002, P_Day4_ = 0.001). In the female-originated LA-N-2 cells, *CHAT* elevation was visible at low concentrations and significant at medium concentrations (Figure 4C). Also, the impact of 440 pM CNTF appeared to peak at or before 48 h, whereas 4.4 nM elicited a dramatic long-term response. Conversely, the male-originated LA-N-5 cells reacted most strongly to CNTF at 440 pM, showing an “inverted U” type dose-response curve (Figure S2). As many as 77.8% of the DE miRs in LA-N-2 reproduced those of a prior sequencing experiment (GSE120520, same layout, only 3 biological replicates).

Apart from this significant overlap, the DE miR profiles showed exclusivities, which might translate to sex-specific regulatory processes in neuronal cholinergic differentiation. However, the log-fold change metric is not ideally suited for assessing the potential impact of expression changes for individual miRs, because it does not reflect mean expression levels. To determine the change in expression, we introduced the count-change metric, a combination of base mean expression and log-fold change, to weigh DE miRs against one another. Count-change values of the 114 miRs detected as DE in both cell lines correlated well, whether the change was observed same- (76 miRs, Spearman’s rho = 0.9066, p < 2.2E-16) or opposite-directionally (38 miRs, rho = −0.9294, p < 2.2E-16) (Figure 4D). Neurokine-induced differentiation of LA-N-2 and LA-N-5 cells further induced a subset of the conserved, primate-specific, and TF-targeting miRs predicted via single cell analysis of cortical cholinergic neurons (Figure 4E, see also Figure 3E). Literature query of two curations of miR precursors in SCZ (Beveridge and Cairns, 2012) and BD (Fries *et al*., 2018) revealed 76 DE miR precursors to be associated with one or both of the diseases (Figure 4F, Data S4). Mature miRs in this case could not be assessed because the curated datasets partly lacked strand information.

### Transcriptional interactions

#### Male and female cells respond to cholinergic differentiation by 5 shared and 12 diverging miR families

miRBase.org currently features 151 human miR families (designated “mir” with lowercase “r”) based on homology; we found members of 71 families DE in LA-N-2 and LA-N-5 cells. Gene set enrichment revealed 5 families which were sex-independently enriched in both cell lines, with highest counts of individual members in the families let-7 (Fisher’s exact test, p = 1.6E-08) and mir-30 (p = 0.015); 12 families were only enriched in one of the two lines, with highest counts in families mir-515 (p = 2.9E-06 in LA-N-2) and mir-154 (p = 3.5E-12 in LA-N-5) (Figure 5A). Of all miR families identified in this analysis, five have previously been associated with SCZ (both sexes: let-7, mir-27; male: mir-181, mir-199; female: mir-10), and three with BD and SCZ (both sexes: mir-30, male: mir-154, female: mir-17).

**Fig. 5.**
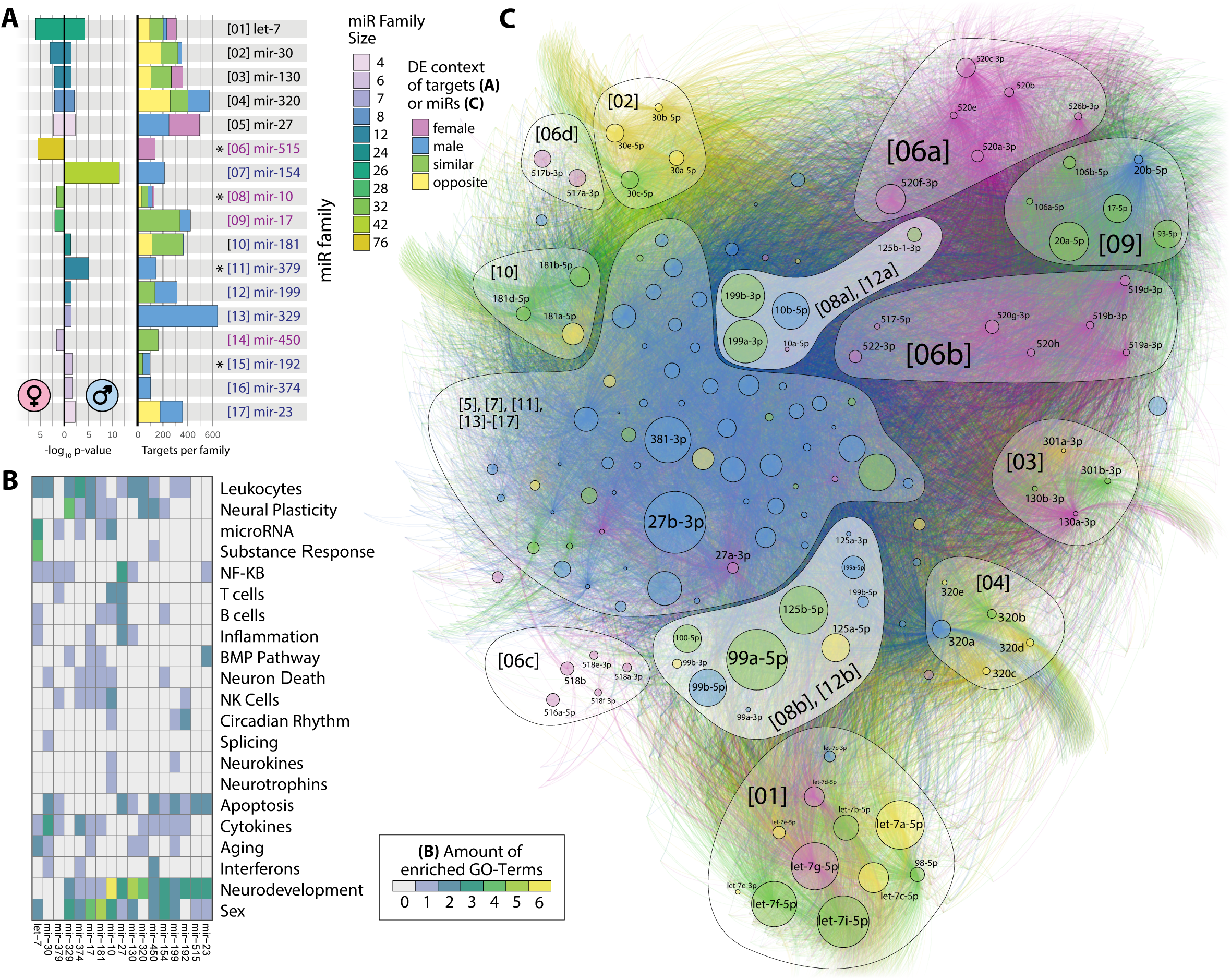
Differentially expressed miR families enriched in LA-N-2 and LA-N-5 cells self-organize in a comprehensive targeting network. **(A)** P-values of 17 enriched families (left) in LA-N-2 and LA-N-5 (female/male symbol) and mean predicted gene target count of each family (right). Color denotes family size (left) or contribution of sex to total target numbers (right) as determined via DE context of individual family members. Asterisks indicate families with significantly smaller gene target sets (see text). **(B)** Gene ontology topics curated for enriched families. Frequent terms are immune-, neurodevelopment-, or sex-related. The mir-10 and mir-199 families show rare association with neurokine signaling and circadian rhythm. **(C)** Comprehensive network of all DE members of enriched families targeting 12495 genes self-separates into family-dependent clusters by application of a force-directed algorithm (46937 unique interactions). miR node size denotes absolute count-change, color denotes DE context. Numbers in brackets correspond to panel (A). mir-10 and mir-199 families form two distinct, sexually dimorphic clusters near the center of the network (lighter background color).

Considering the mechanism of action of miRs, i.e. their multiple-targeting behavior, it is of interest how broadly these families act on gene targets. Seeking the per-family mean target count (via *miRNet*), we found that families enriched in only male or female cells had significantly less targets than those enriched in both cells (217 vs. 378, Welch two-sample t-test, p = 0.001). Relative to their size, 4 families show significantly lower target numbers than all other families: mir-10 (p = 0.016), mir-192 (p = 0.042), mir-379 (p = 0.011), and mir-515 (p < 0.001) (Figure 5A, right hand side). This might indicate a spectrum of functional categories, from broadly acting families such as let-7 with sex-independent function to families with narrow target profile such as mir-10, whose restricted function can associate with sex-specific impact.

#### Enriched families regulate genes involved in immunity, neuronal development and sex

To assess their putative functions, we performed GO enrichment analysis of the 300 most-targeted genes in each of the enriched miR families. This revealed involvement in 1124 biological processes, manual curation of which enabled us to infer the functional roles of each family (Figure 5B, Data S5). Resultant terms related to neuronal development, cytokine- and leukocyte-mediated immune processes, and sexual dimorphism. Compatible with the hypothesis of “broad to narrow” functioning families, we also found terms with lower incidence and higher specialization in subgroups of families.

#### Differentially expressed families self-organize in a force-directed network

Unbiased graphical network analysis of the 212 DE miRs from all enriched families and their 12495 targeted genes (via *miRNet*) yielded a complex interactome. A force-directed algorithm (Jacomy *et al*., 2014) yielded apparent clustering and, in some cases, subdivisions of families (Figure 5C). Two main clusters emerged for male and female DE miRs, from families mir-515 and mir-17 (female) and families mir-154, −379, −329, −129, −374, and −23 (male). Most other families show marginal localization, distributed around the network’s edge. Three small families show high centrality and high count-change: mir-27, targeting particularly many genes, and the neurokine-associated mir-10 and mir-199, with lower target count.

#### The mir-10 and mir-199 families show sexual dimorphism of expression and ontological association to neurokine and circadian mechanisms

Single members of the mir-10 and mir-199 families were detected in each of the DE categories: female only, male only, and similar-as well as opposite-directionally. In the comprehensive network, mir-10 and mir-199 form two distinct clusters (“[08a], [12a]” and “[08b], [12b]”) composed of opposite strands of their respective miR precursors. Ontologically, they associate with neurokine and circadian genes. Further, 125a/b-5p (members of mir-10), and 199a/b-5p belong to those miRs predicted to target genes expressed in single cholinergic neurons (see Figure 3E/4E).

Further analysis requires a closer look into subnetworks (e.g. of individual families), which are astoundingly heterogeneous in size, layout, and in their individual sexual dimorphism; a comprehensive assessment is outside the scope of this study. Fully interactive versions of all individual enriched family networks and further information can be accessed at https://slobentanzer.github.io/cholinergic-neurokine; the entire network in tabular format is provided as a resource (Data S6). Below, we describe the cholinergic/neurokine subnetwork.

### The cholinergic/neurokine interface

#### Creation of a cholinergic/neurokine sub-connectome by gene- and miR-filtering

To exemplify the proposed complexity reduction technique, we looked for the common denominator enclosing all aspects of this study; a limited connectome analysis requires a defined set of genes and miRs. In the beginning, we performed an unbiased analysis of sexual dimorphism in SCZ and BD, which implicated processes of neuronal, immunological, and circadian origin (Figure 1). Since our experimental data is based on cholinergic processes, we compiled a list of relevant cholinergic genes (Soreq, 2015), adding to it genes from pathways that had emerged in the previous analyses: neurokine signaling and circadian rhythm. Returning to the collection of web-available patient data, we subjected this limited set of 76 genes and their 18 neuronal TFs to differential expression analysis (Data S7).

The miRs were gradually filtered by multiple consecutive steps: (*i*) Permutation analysis of comprehensive miR targeting data specific for genes expressed in cholinergic neurons (Figure 3) yielded a list of miR candidates that shows overlap with (*ii*) miRs DE in our two models of neurokine-induced cholinergic differentiation (Figure 4). (*iii*) We included only families of miRs we found to be enriched in differential expression (Figure 5). This filtering process yielded 69 miRs from 12 families (Data S8), which were assembled in a force-directed network with the 94 genes of the previously compiled list; as a “spike-in”, we added miR-132-3p (DE in LA-N-5), a well-studied miR which is known to influence cholinergic processes (Shaked *et al*., 2009; Shaltiel *et al*., 2013; Hanin *et al*., 2018). The resultant network (Figure 6A) shows high structural homology to the comprehensive Figure 5C network, with similar groupings and spatial organization of families.

**Fig. 6.**
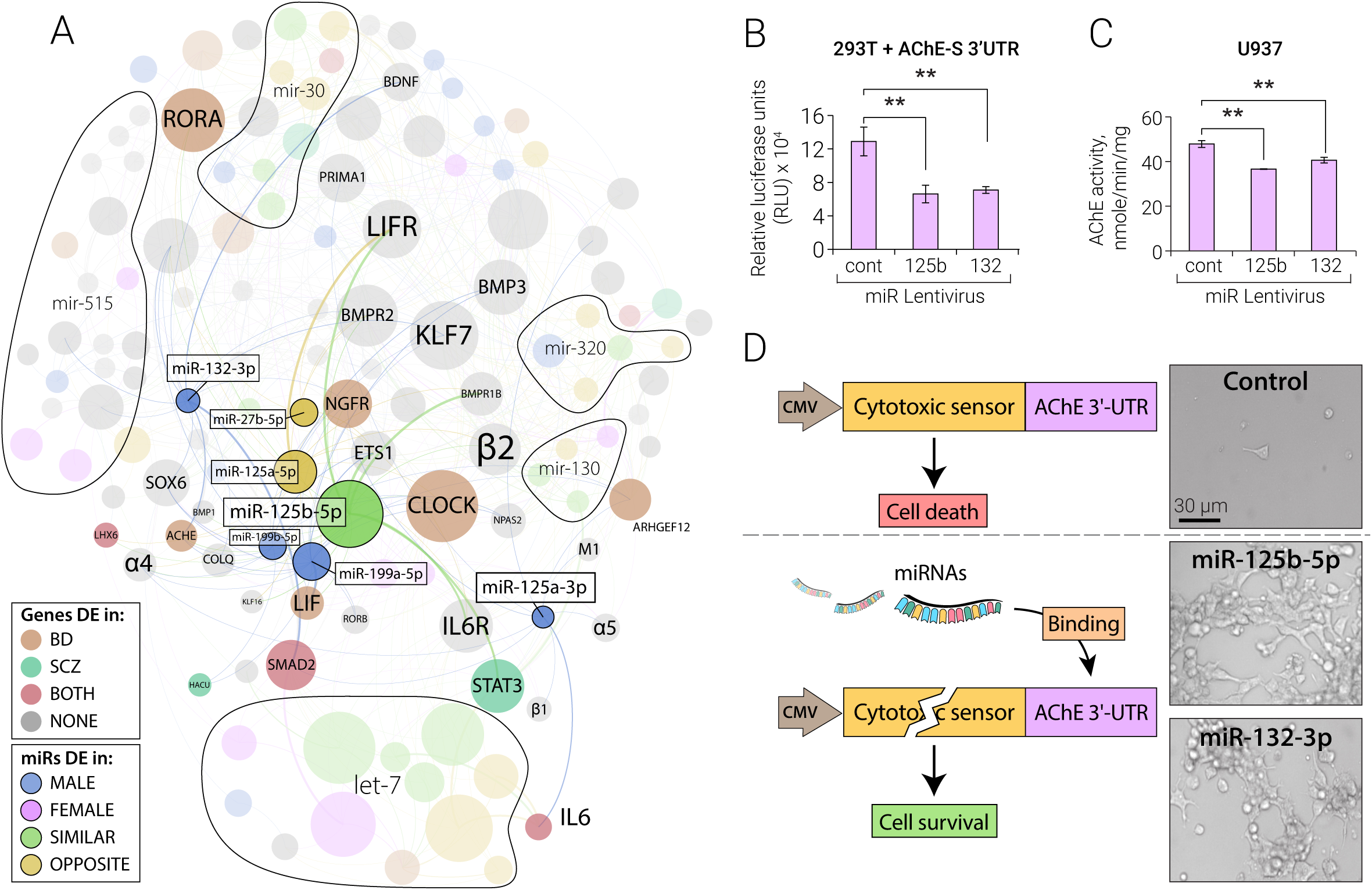
The cholinergic/neurokine interface and experimental validation of AChE targeting by hsa-miR-125b. (**A**) miR families mir-10 and mir-199 pose a sexually dimorphic interface of cholinergic, neurokine, and circadian regulation by targeting nicotinic/muscarinic (e.g. *α4β2, M1*) and neurokine receptors, transcriptional regulators of cholinergic differentiation (*LHX, STAT*) and circadian rhythm (*CLOCK, RORA*), AChE and the AChE linker proteins *PRIMA1/COLQ*, and high affinity choline uptake (*HACU*). Members of mir-10/199 families, “spike-in” miR-132-3p, and their targeted genes are shown in color, other miR families that passed the multiple filtering are indicated as areas. miR node size corresponds to count-change, gene node size to connectivity, color and thicker edges indicate DE context and experimentally validated connections. (**B-D**) Validation experiments of AChE targeting by miR-125b-5p, with miR-132-3p as a positive control. (**B**) Lentiviral expression of miRs-132 and −125b suppresses luciferase fused to the 3’-UTR of AChE in HEK293T cells. (**C**) Lentiviral expression of miR-132 and miR-125b suppresses the endogenous AChE hydrolytic activity of U937 cells with similar efficacy. (**D**)Life/death assay of stably transfected HEK293T cells carrying the AChE 3’-UTR fused to a cytotoxic sensor, and co-transfected with miR-125b-5p, −132-3p, or control plasmids. Cells survive in case of binding of miR-132-3p and −125-5p to the 3’-UTR.

#### mir-10/199 family members are pivotal factors in the cholinergic/neurokine interface

Of the 23 genes we found perturbed in the re-analyzed SCZ/BD patient data, miR-125a-5p and miR-125b-5p targeted 9 genes each, followed by let-7a-3p, let-7f-1-3p, miR-199a-5p, miR-199b-5p, and miR-30a-5p (targeting 7 genes each). In this sub-connectome, the most-targeted DE genes (as indicated by node size) are the circadian regulators CLOCK and RORA, and the neurokine pathway genes LIFR and STAT3. Members of mir-10/199 are intricately involved in the control of all of these factors, indicated by their closeness and central location.

#### mir-10 family member miR-125b-5p is highly differentially expressed in LA-N-2 and LA-N-5 and targets AChE

Of all miRs in the reduced set, hsa-miR-125b-5p exhibits the highest absolute count-change, and displays most experimentally validated targeting relationships with cholinergic/neurokine genes, targeting several inflammation-related neurokine pathway genes (miRTarBase accessions; IL6: MIRT022105, IL6R: MIRT006844, JAK2: MIRT734987, LIF: MIRT001037, LIFR: MIRT732494, STAT3: MIRT005006), and other inflammatory pathways (e.g. TNF: MIRT733472, IRF4: MIRT004534). Additionally, hsa-miR-125b-5p harbors a seed sequence predicting both subunits of the nicotinic α4β2 receptor and AChE as targets.

To experimentally test the capacity of hsa-miR-125b-5p to impact cholinergic signaling, we performed a cell culture luciferase assay, cell death and endogenous enzyme suppression tests on the secreted AChE protein (Figure 6 B-D). This analysis revealed functional suppression of AChE by miR-125b-5p with similar efficacy to that of the positive control miR-132-3p (Shaked *et al*., 2009; Hanin *et al*., 2018). Lentiviral infection with miR-132 and −125b led to equally efficient suppression of luciferase activity in HEK293T cells stably transfected with a plasmid expressing luciferase fused to the AChE mRNA 3’-UTR (one-way ANOVA: p = 0.004, Figure 6B). Further, lentiviral infection of human monocyte-like U937 cells with miR-125b-5p suppressed endogenous AChE hydrolytic activity (one-way ANOVA: p = 0.005, Figure 6C), and co-transfection of miR-125b-5p with a lentiviral vector expressing a cytotoxic sensor fused to AChE 3’-UTR resulted in cell survival with similar efficacy to that of the positive miR-132-3p control in HEK293T cells (n = 3, Figure 6D). In addition to the already experimentally validated interactions, this makes hsa-miR-125b-5p a prime candidate of cholinergic/neurokine mediation.

## Discussion

The magnitude of interactions shown by network analyses, further complicated by high between-tissue variability, presents one of today’s largest obstacles in microRNA research. As miRs exert individual effects of low impact, the outcome of their cooperative action can be more adequately represented by a network approach than by traditional molecular interaction studies (Salta and De Strooper, 2017). While experimental validations are necessary, they often significantly exceed the scope of scientific publications even for one single miR. On the other hand, purely bioinformatical interaction studies lack predictive power, particularly for non-canonical miR targeting, with high false positive and false negative rates (Hart *et al*., 2018). Bioinformatically supported high-throughput techniques such as short RNA sequencing can serve as an integrative middle ground. Our study addresses this purpose by facilitating the identification of manageable numbers of interaction partners for deeper analyses of subnetworks involved in specific ontological categories of coding genes. We chose the cholinergic/neurokine interface as an example due to our interest in cholinergic transmission, which is intrinsically linked to many processes relevant in SCZ/BD (Figure 2). Co-expression of cholinergic and neurokine markers in single cortical cells (Figure 3), the pro-cholinergic influence of neurokines on our model cell system (Figure 4), and the identification of neurokine signalling and circadian rhythm in GO enrichment analysis of specific miR families (Figure 5) highlight the relevance of this choice to our current study.

### Limitations

We chose a human cellular model over *in vivo*-experimentation for several reasons. First, implementation of cholinergic differentiation in the brain of a living animal is not straightforward, and individual types of cholinergic neurons are quantitatively inferior to supporting cells in the cortex (von Engelhardt *et al*., 2007). Second, a recent study (Naqvi *et al*., 2019) demonstrates that most sex bias in gene expression has arisen since the last common ancestor of boroeutherian mammals (including mouse, rat, dog, macaque, and human), resulting in inadequate representation of sexual dimorphism in any non-human model organism; and third, current animal models of SCZ and BD do not faithfully represent human pathology, showing no predictive power for clinical efficacy of therapeutics (Jones *et al*., 2011). Additionally, the resolution of differential expression analysis in sequencing is much higher in a homogeneous cell population such as neuronal cell culture, as sexual dimorphism can also manifest in distinct tissue composition (Naqvi *et al*., 2019). However, this choice also entails limitations: Although very similar, LA-N-2 and LA-N-5 are immortalized cells derived from two distinct donating individuals, which must be considered when interpreting sexual dimorphisms based on their total transcriptional divergence. Specifically, the stronger response of LA-N-5 to CNTF might have influenced the detection limit of miRs designated “male” by their lack of detection in LA-N-2, partly leading to the dominant male cluster in the comprehensive network. On the other hand, this increases the power of detection of “female”-designated miRs (e.g. mir-515), because they were not detected in LA-N-5 cells in spite of higher sensitivity.

Notwithstanding these limitations, we consider our approach a viable alternative in situations which require a sophisticated perspective on sexual dimorphisms, particularly in diseases where representative animal models are not available. In the present study, we defined ontological categories implicated in SCZ/BD dimorphisms via meta-analysis of deposited patient data, ascertained co-expression of our principal systems of interest in the context of single cortical cells, assessed differential expression via suitable cellular models, and comprehensively analyzed the transcriptional interactions of identified miRs in their genome-scale regulatory network. We used, as well as created, comprehensive web-available resources that can aid focused studies by generating hypotheses and candidate lists, or by putting experimental results into context. We advocate the application of this integrative methodology to make use of the many excellent data collections created by modern science.

### The cholinergic/neurokine interface

Our approach identified a trophic role for neurokine signalling in cortical cholinergic systems, particularly for IL-6, distinguishing these cortical neurons from the NGF-dependent basal forebrain population. The consistent identification of inflammatory processes across all aspects of our integrative study protocol lends further support to the neuroinflammatory aspect of SCZ/BD, for example via IL6-mediated temporary overstimulation (Lurie, 2018). Basal neurokine levels in the brain are comparatively low, and current methods of measurement do not allow an analysis of short-term fluctuations in living human subjects. However, it has been shown that CNTF can influence behaviour through its impact on cholinergic neurons in the arcuate nucleus (Couvreur *et al*., 2012). Paracrine control of cholinergic neurons by neurokine-affected supporting cells such as glia or astrocytes is also physiologically feasible, pending further analyses.

Compatible with our previous findings (Cohen *et al*., 2002), we found a paramount role of *CLOCK* in the mir-10/mir-199 regulatory network, indicating circadian regulation of neurokine control over cholinergic signalling. In RNA sequencing of 600 prefrontal cortices of SCZ patients and controls, *IL6ST* (aka gp130), and the host gene for the *IL6ST*-targeting hsa-miR-335, were among the top 50 imprinted genes (Gulyas-Kovacs *et al*., 2018). While the data used in that study yields surprisingly little DE genes, it discovered significant depression of the *CNTFR* and cholinergic receptors, accompanied by an elevation of *CLOCK* and *RORA*. Conversely, a knockdown of *CLOCK* in *in-vitro* differentiated human neurons (Fontenot *et al*., 2017) caused parallel neurokine receptor and cholinergic transcript perturbations, suggesting an intrinsic relationship of these systems.

### miRNA families

Our tests also implicated several novel factors pivotal for control of cholinergic function in BD and SCZ pathology. Recently, SCZ and BD have come to be recognized as instances of a spectrum of transcriptional perturbations with increasing “transcriptomic severity” from BD to SCZ (Gandal *et al*., 2018). Our approach identified 3 miR families associated with both diseases and 5 associated with SCZ, but none were associated with BD alone, retracing the aspect of increasing severity from a non-coding perspective. The closely related mir-10 and mir-199 families are intrinsically sexually dimorphic, have previously been associated with SCZ (Beveridge and Cairns, 2012; Szatkiewicz *et al*., 2014) and show a comparatively small number of gene targets, predicting a focused regulatory role. While individual members of large families such as let-7 or mir-515 often show distinct individual target profiles, mir-10 family members demonstrate overlapping functional features of their targets, even between their −5p and −3p variants. The evolutionarily conserved miR-132-3p, through its suppression of AChE, interrupts the cholinergic modulation of acute stress (Shaltiel *et al*., 2013), immune response (Shaked *et al*., 2009) and metabolic activities (Hanin *et al*., 2018) and leads to a transient increase of IL-6, which can be mitigated by nicotine (Shaked *et al*., 2009). However, miR-125b-5p but not miR-132-3p may also target the α4β2 nicotinic receptor subunits, such that its modulation could have an additional impact. Additionally, miR-125b-5p can influence inflammatory processes directly by targeting 5-lipoxygenase (Busch *et al*., 2015), and indirectly via regulation of epigenetic controllers (Zhang *et al*., 2017). Moreover, miR-125b-5p is directly induced by the Vitamin D-receptor (Giangreco *et al*., 2013), and decreased Vitamin D levels are thought to be a risk factor for SCZ/BD development (Cieslak *et al*., 2014). miR-125-5p hence possesses several cholinergic and non-cholinergic features potentially relevant to SCZ/BD pathology.

Our experiments also indicate other miRs as potential mediators of sexually dimorphic cholinergic, neurokine, and circadian functions of human neurons. For example, miR-124 is DE between LA-N-2 and LA-N-5 after neurokine induction. A recent analysis of a complete miR-124 knockout in human iPSCs (Kutsche *et al*., 2018) found extensive subsequent transcriptional perturbations, in STAT5B among others, and a shift from glutamate to ACh in the resulting functional neurons. Although the authors did not address this cholinergic aspect, their data and ours co-indicate an influence of miR-124 on the neuronal cholinergic phenotype.

### Therapy

At present, treating cognitive malfunctioning through stimulation of the cholinergic system is limited to AChE inhibitors and nicotinic/muscarinic agonists (Rowe *et al*., 2015). Stimulations of nicotinic (e.g. α4β2, the α5-subunit, and α7 (Koukouli *et al*., 2017)) and muscarinic (e.g. M1) receptors (Vijayraghavan *et al*., 2018) display a therapy-limiting lack of specificity for individual receptors/subunits. On the other hand, neurokine-based interventions have repeatedly failed due to intolerable side effects, even when applied directly to the brain (Mufson *et al*., 2009). In comparison, antisense oligonucleotide therapeutics can simultaneously target multiple disease-relevant genes, be it as mimic/antagonist of an existing miR, or a synthetic oligonucleotide aptamer engineered for a specific target profile. The success of oligonucleotide therapy will depend on calibrating its impact on target genes and achieving a positive balance between target and off-target effects (Greenberg and Soreq, 2014). Therapy with miR-like molecules can also potentially ameliorate tachyphylactic effects by targeting a gene and its regulatory elements at the same time (essentially via feed-forward-loops (Guzzi *et al*., 2015)), as implicated by the nested regulatory circuits we observed. Identification of lead molecules for development of therapeutics with defined profiles largely depends on an integrative approach combining gene- and TF-targeting.

Recent clinical advances in personalized medicine involve evaluating the benefit of sex-specific therapeutic adjustments in cholinergic medication, at least for Alzheimer’s disease (reviewed in (Giacobini and Pepeu, 2018)), but the molecular examination of cholinergic systems (e.g., with respect to co-expression of sex hormone receptors) has so far been limited to the basal forebrain Ch1-4 nuclei. While some of the datasets covered in the present study allow sex-specific analyses, many questions concerning sexual dimorphisms remain unaddressed and warrant extensive future studies. The advent of single-cell sequencing will hopefully enable application of this methodology in actual patient tissues. More immediately, our study paves the way for the investigation of other processes by application of our method to other ontological categories, e.g. dopaminergic signalling or innate immunity. High quality datasets large enough to allow statistical analyses can further enable the extension of this approach to different brain regions and other psychiatric and non-psychiatric polygenic disorders such as sporadic Alzheimer’s and Parkinson’s diseases.

## Acknowledgments

The authors are grateful to in-depth comments by Drs. Naomi Habib, Jerusalem and Andreas Meisel, Berlin and to Dr. Ester Bennett, Jerusalem for assistance with the RNA-sequencing.

## Author contributions

Conceptualization, H.S. and S.L.; Methodology, S.L.; Software, S.L.; Investigation, S.L. and G.H.; Resources, J.K. and H.S.; Data Curation - S.L.; Writing - Original Draft, S.L.; Writing - Review and Editing, S.L., J.K. and H.S.; Visualization, S.L.; Supervision, J.K. and H.S.; Project Administration, J.K. and H.S.; Funding Acquisition, J.K. and H.S.

## EXPERIMENTAL MODEL AND SUBJECT DETAILS

### Cell lines

We employed LA-N-2 (female) (DSMZ Cat# ACC-671, RRID:CVCL_1829) and LA-N-5 (male) (DSMZ Cat# ACC-673, RRID:CVCL_0389) cells as our main experimental model system (McManaman and Crawford, 1991). The cells were purchased at DSMZ (Braunschweig, Germany). These cells respond to differentiation by several neurokines (CNTF, LIF, IL-6) by cholinergic differentiation corresponding with elevation of choline acetyltransferase (ChAT), the central cholinergic marker (mRNA, protein, and activity) as well as its intronic vesicular ACh-transporter gene, vAChT (aka SLC18A3). Cells were maintained at 37°C in 8% CO_2_ atmosphere in medium consisting of 1:1 DMEM and RPMI 1640, with 20% FCS, with weekly splits. Experiments were performed between splits 2-6 after thawing.

HEK293T and U937 cells were purchased at ATCC and maintained according to ATCC guidelines. Experiments were performed between splits 2-10.

### Human patient data

We used previously published data sets for several analyses. These comprise several cortical data sets in raw format (Affymetrix) from NCBI GEO: GSE35978 (Chen *et al*., 2013) (SCZ & BD), GSE53987 (Iwamoto *et al*., 2005) (SCZ & BD), GSE12649 (Lanz *et al*., 2015) (SCZ & BD), GSE17612 (Maycox *et al*., 2009) (SCZ), GSE21138 (Narayan *et al*., 2008) (SCZ), GSE5392 (Ryan *et al*., 2006) (BD); next-generation sequencing from NCBI GEO: GSE80655 (Ramaker *et al*., 2017), GSE106589 (Hoffman *et al*., 2017), GSE68559 (Webb *et al*., 2015), GSE96659 (Fontenot *et al*., 2017), GSE45642 (Li *et al*., 2013); data of DLPFC sequencing of 600 SCZ patients and controls was obtained from the Common Mind Consortium (http://www.synapse.org/CMC).

## METHOD DETAILS

### Methodological outline

Focusing on a well-defined set of genes aims to avoid some of the data loss that occurs when patient tissues comprising multiple cell types are homogenized for analysis. This is particularly relevant for cortical cholinergic interneurons, where the cell type of interest is numerically inferior, and transcriptomic data can be “diluted” by the other, more prominent cell types, such as glia or astrocytes. The consequent increase in detection threshold can be re-lowered by reducing the number of genes tested. Analysis of single cell datasets can ascertain that the genes analysed are representative and fairly unique to the cell type in question, as was the case for cholinergic markers in our study, and utmost care should be taken for any gene set of interest and any kind of tissue subjected to this kind of analysis.

Our approach is based on the availability of suitable amounts of patient data in web-available form, combined with a standardized pipeline of statistical pre-processing to equilibrate individual statistical influences. Regardless of how patient data and the subset of genes of interest are selected, the reduction in number of analyzed genes has to be performed after application of the linear model to the batch- and covariate-corrected data to avoid interference with correct linear regression. Once the expression data (be it array- or sequencing-derived) has been corrected for batch effects and covariates, and outliers have been removed, it can be analyzed with a suitable differential gene expression algorithm. The resulting, individual datasets should ideally converge on similar logFC values, but may also show controversy between individual experiments, which can result from a multitude of factors from biological variety to sampling procedures or exact tissue composition, all of which have to be interpreted at the discretion of the scientist.

### Neuronal differentiation/short RNA sequencing

#### Model

LA-N-2 and LA-N-5 cells respond to ciliary neurotrophic factor (CNTF) by cholinergic differentiation corresponding with elevation of choline acetyltransferase (ChAT), the central cholinergic marker (mRNA, protein, and activity) as well as its intronic vesicular ACh-transporter gene, vAChT (aka SLC18A3). We measured the changes in short RNA levels following this intervention at several time points.

#### Differentiation

To determine effective concentrations of the differentiation agent, we performed a dose-response experiment with both cell lines. Cells were seeded at approximately 200 000 cells per well in 12-well plates, and after 24h incubated with 1, 10, or 100 ng/ml CNTF (Sigma-Aldrich). Dose-response was measured by qPCR of CHAT mRNA, at time points 30 minutes, 60 minutes, 2 days, and 4 days. For each sample, a corresponding control culture was generated.

#### RNA extraction for qPCR and sequencing

RNA was extracted in biological quadruplicates using TRIzol according to the manufacturer’s instructions, as described in (Hanin *et al*., 2018). RNA was precipitated using ethanol, washed, and air dried before resuspension in RNAse-free water. Concentration was measured by Nanodrop 2000 (ThermoFisher Scientific), RNA quality was measured via Bioanalyzer 2100 (Agilent). RNA quality for all samples was near optimal (RIN > 9).

#### Quantitative real-time PCR

RNA was analyzed on a BioRad CFX96 real-time PCR cycler using PowerUp SYBR Green Master Mix (Applied Biosystems) in technical duplicates. Primers were designed using primer3 and are as follows (5’ to 3’): ChAT [FW: CAC TTG GTG TCT GAG CA, RV: AGT TTC TGC TGC AGG GTC TC], ACTB (housekeeping) [FW: GCT GTA TTC CCC TCC ATC GT, RV: CTT CTC CAT GTC GTC CCA GT]; additional primers were ordered from BioRad, Germany: vAChT [PrimePCR “qHsaCED0047922”], RPLP0 (housekeeping) [PrimePCR “qHsaCED0038653”]. Data were analyzed with BioRad CFX manager and expression values (normalized to housekeeping genes) exported for statistical testing in R.

#### Short RNA sequencing

Short RNA sequencing was performed using Illumina NextSeq 550 according to the manufacturer’s instructions, after cDNA library preparation using the NEBNext Multiplex Small RNA Library Prep Set for Illumina (New England BioLabs) as described (Bekenstein *et al*., 2017). Sequenced reads were aligned to miRBase v21 sequences via miRExpress (Wang *et al*., 2009) (version 2.1.4). Differential expression was determined via R/DESeq2 (Love *et al*., 2014).

#### The count-change metric

We calculated the count-change for individual miRs by combining base mean expression with the de-logarithmized fold-change (from DESeq2 output).

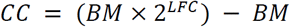

CC: countChange, BM: baseMean, LFC: log2-fold change

It is important to note that the count-change metric, by deriving from the base mean expression across samples, is dependent on sequencing depth, and thus is not instantly generalizable, for instance when comparing different experiments. However, it could be normalized to a degree by considering the total amount of raw reads generated from each sample.

### Whole transcriptome meta-analysis

#### Data preparation

We processed web-available patient transcriptome data sets by state-of-the-art procedures, analogous to Gandal et al. (Gandal *et al*., 2018), to generate transcriptional disease profiles. The following R packages (Bioconductor) were used according to the developer’s instructions: Raw data read, RMA normalization, RNA degradation: affy (Gautier *et al*., 2004). Batch correction (as per chip scan date) involved: sva (Combat) (Johnson *et al*., 2007). Outliers were removed as described in (Gandal *et al*., 2018). The array probes were annotated via ENSEMBL gene ID, database version v75, to ensure congruency with prior analyses, using biomaRt (Durinck *et al*., 2009), and were collapsed via the collapseRows function of WGCNA (Langfelder and Horvath, 2008). Prior to application of the generalized linear mixed model (nlme (Lindstrom and Bates, 1990)), datasets were rebalanced and regressed to correct for technical and biological covariate influences.

#### Regression analysis

Meta-analysis on all genes present in all datasets (12,391 genes in total) employed a generalized linear mixed model to account for variation per dataset and individual, yielding log_2_-fold change (logFC) values for each gene between control and disease, which were correlated between SCZ and BD using Spearman’s method (Spearman, 1904). To determine significance, the meta-analysis process was repeated 10,000 times with randomized case/control status, forming a permutation null distribution of the individual correlation coefficients (Figure S3). Of note, several cholinergic and neurokine genes were missing from the whole-genome meta-analysis because of annotation deficits (*CHRNA7, CHRM1, LHX8, CHKB, PRIMA1, CNTF*), and, at this stage, could not be easily re-introduced.

#### Gene ontology (GO) enrichment analyses

To focus on the differences, as opposed to the similarities, between BD and SCZ patient brain transcriptomes, we performed GO enrichment analysis on the genes showing the highest rank differences between datasets (Figure S4), using the R package topGO (Alexa *et al*., 2006). Briefly, we evaluated (smaller) groups of genes for enrichment against a (bigger) background of genes for presence in individual GO terms exceeding statistical estimates. The background comprised of the first 2000 genes according to the applied rank system, computed as a function of Spearman’s rank differences of regression beta values (logFC) between the two compared groups, either as absolute values or “as is” (elevation in one group as opposed to the other). The target genes of the analysis were defined as the 100 top ranked genes (top 5% of background), unless otherwise stated. Statistically significant GO results were compiled and curated for CNS-relevant terms.

#### Sex influence on transcriptomic differences

Studying web-available datasets of non-degenerative mental disease patients revealed that SCZ and BD datasets possessed sufficient numbers to allow sex-discriminative meta-analysis of statistical significance. Hence, we repeated the above steps for the individual subsets of male and female patients, with the sole change of eliminating the covariate regression for sex, as this would preclude further analysis of this variable, in 4 distinct GO enrichment group comparisons between SCZ-biased, BD-biased, male-biased, and female-biased genes.

### miR-gene-TF-targeting: miRNet

To address miR-mRNA targeting relations, we developed an integrated miR-targeting graph database (*‘miRNet’*) out of publicly available validated and predicted data, implementing a scoring system derived from 10 leading prediction algorithms (Dweep and Gretz, 2015) based on their statistical performance at the whole-genome level. To facilitate the selection process, targeting data was cumulated by summing the amount of positive “hits” of prediction algorithms (1 point) and positive experimental validations (only “strong” evidence, miRTarBase, 10,5 points) of the targeting relationship, yielding a targeting score between 0 and 20,5. We found a minimal score of 6 to suffice for a good balance between type I and II errors. In case of whole-genome interactions, this threshold was raised to 7 to enable computational accessibility in all steps (in this case, graphical analysis was the bottleneck). The database was supplemented by comprehensive transcription factor targeting data via bioinformatically processed “cap analysis of gene expression” (CAGE (Hon *et al*., 2017)), focusing on brain tissues (Marbach *et al*., 2016), including the tissue-specific transcriptional activities (Figure S5). The code used to create and test this database is available in the accompanying repository. A public release is planned, but not completed at the moment; requests can be directed at the corresponding author.

Analyses performed using this database were implemented in R using the ‘RNeo4j’ package.

#### Limitations

Through recent methodical and bioinformatical advances, targeting data of transcription factors and miRs has become comprehensive. However, these “complete” datasets are subject to limitations derived from the methods used to accumulate raw data. For TF-targeting, CAGE 5’ peaks were analyzed towards their correlation with gene expression in all available tissues. A cholinergic example of limitations derived from this data involves the *CHAT* gene, which does not provide a measurable CAGE peak, leading to non-representation in the targeting dataset. In those cases, targeting data acquired through conventional methods have to be substituted, but this cannot be easily done for all affected genes, and is also not comprehensive. In the case of miR-targeting, validated interactions are based on experimental work mainly performed on rodents, leading to a research bias towards evolutionarily conserved miRs. Primate-specific miRs therefore are under-represented, at least in validated data.

#### Permutation targeting analyses

In the current state of comprehensive data on miR-gene (and TF-gene) targeting, no statements can be made with absolute certainty. Thus, an approach which considers relative measures is preferable. For this reason, whenever whole-genome/whole-miRnome targeting was concerned, we employed random permutation of the prediction dataset or each single predicted miR against a randomized background of the same size as the original set (also considering family- and precursor-relationships). The resulting null distribution yields a basis for determination of a false discovery rate.

#### Neurokine-induced miRs and the cholinergic/neurokine pathway

Gene targets of the 490 differentially regulated miRs following CNTF exposure were determined by *miRNet* query (targeting score minimum of 6). The full network, originally comprising ∼160,000 unique relationships, was re-filtered by raising the threshold to score minimum of 7 to be computationally accessible. The resulting network and individual miR-family-subnetworks were plotted using a force-directed layout (Force Atlas 2) in gephi.

#### Single cell sequencing data set analysis and permutation

We analyzed 4 web-available datasets of brain single cell gene expression (Darmanis *et al*., 2015; Zeisel *et al*., 2015; Habib *et al*., 2016; Tasic *et al*., 2016) for neurokine signaling transcripts in cholinergic neurons, identified by their expression of the ACh-synthesizing enzyme choline acetyltransferase (ChAT), and its embedded gene encoding the vesicular acetylcholine transporter SLC18A3, also known as vAChT. Raw gene expression data was normalized and then clustered and plotted via R/pheatmap (Kolde and Kolde, 2015). The genes expressed in more than one sample per dataset were enriched for targeting of TFs and miRs by random permutation analysis (via *miRNet*), with 10 000 permutations for conserved and primate miR-target relationships, and human CNS TF interactions.

### Subset analyses

#### Cholinergic genes, transcription factor and neurokine analyses

To follow our cholinergic interest, we used a recent review (Soreq, 2015) to define a core set of genes, adding to it the neurokine and circadian pathway genes indicated in the previous analyses (to a total of 76 genes). Via *miRNet* permutation (Data S2), we identified 18 brain-expressed “cholinergic” TFs (p < 0.05) and their CNS transcriptional activity towards each targeted cholinergic gene. These 94 genes (Data S7) were then subjected to differential expression analysis in the deposited patient datasets.

#### Execution of subset analyses

To separately analyze male and female data in the original datasets, we repeated the sex-independent analyses of the identified 94 cholinergic genes and TFs using the limma (Ritchie *et al*., 2015) pipeline. Pre-processing was identical to the whole-transcriptome approach, and dataset reduction involved restricting the output table (*topTable()* function) to the studied genes. This further allowed controlling of missing genes of interest by manually solving problems of annotation, which in a whole-genome analysis would have led to loss of information on, e.g., the nicotinic α7 and stress-responding M1 cholinergic receptors.

### miR-125b-5p validation

To test binding of hsa-miR-125b-5p to the acetylcholinesterase (AChE) 3’-UTR, we performed vector-based assays via suppression of a cytotoxic sensor as well as of Renilla luciferase, both fused to the AChE-3’-UTR. Briefly, the 3’ untranslated region (3’-UTR) of human AChE mRNA (Soreq *et al*., 1990) was cloned into the microRNA Target Selection System plasmid (System Biosciences, CA, USA) multiple cloning site, using EcoRI and NotI restriction enzymes (New England Biolabs). All plasmids were verified by DNA sequencing. For luciferase assays, HEK293T cells were transfected with miRNA Target Selection-AChE-3’UTR, and selected in the presence of Puromycin for 3 weeks. Stably transfected HEK293T (293T-AChE 3’UTR) cells were grown on 12-well plates and infected with lentiviruses expressing miR-125b-5p, miR-132-3p or a negative control sequence. After 48 hours incubation, cells were analyzed using the Dual Luciferase Assay kit (Promega, WI USA) and Luciferase activity was measured using an Envision luminescent plate reader (Perkin-Elmer, Waltham, MA), essentially as previously described (Hanin *et al*., 2014). For each reporter construct, renilla luciferase activity was normalized according to that of the firefly. Normalized activity after infection of miR-132-3p or miR-125b-5p was expressed as relative to that obtained after infection with the same plasmid with miR negative control. For life/death assay, a similar protocol was used. Stably transfected HEK293T (293T-AChE 3’UTR) cells were infected with lentiviruses expressing miR-125b-5p, miR-132-3p or a negative control sequence. 72 hours post-infection a cytotoxic reporter fused to AChE 3’-UTR was added to the media and cells were kept for an additional 5 days to assess their viability. Statistical significance was determined using ANOVA with correction for multiple testing. To show effects of changes in this miR’s levels on real-life protein activities, we performed an AChE hydrolytic activity assay following infection of human monocyte-like U937 cells with hsa-miR-125b-5p, miR-132-3p or a negative control lentiviral vector. AChE hydrolytic activity levels were assessed by kinetic measurements of the hydrolysis rates of 1 mM acetylthiocholine (ATCh, Sigma) at room temperature, following 20 min incubation with and without 5×10^−5^ M tetraisopropyl pyrophosphoramide (iso-OMPA, Sigma), a specific inhibitor of butyrylcholinesterase, to selectively assay for AChE-specific or total cholinesterase activity. Each sample was assayed in at least 3 biological replicates. In all cases, hsa-miR-132-3p served as a positive control.

## QUANTIFICATION AND STATISTICAL ANALYSIS

Statistical analyses were performed in R. Small sets of continuous variables, such as qPCR and miR-125b-5p validation experiments, were tested using Welch’s Two Sample t-test (because in most cases, equal variance could not be assumed; R/t.test) and ANOVA; statistical significance was assumed at p < 0.05. For sequencing count data, the negative binomial generalized linear model was tested using the R/DESeq2 package (“Wald” test including log-fold change shrinkage) to detect differentially expressed miRs, using the supplied independent filtering and correction for multiple testing at an alpha level of 0.1. In cases where additional power was desirable or p-values could not be obtained by other means (transcriptome meta-analysis, whole genome miR-targeting), permutation analysis was performed. This comprised random assignment of test variables in the same size as the original test set and repeating the analysis for a large number of times, such that a null distribution of values could be generated, which can be used to determine a false discovery ratio for the original result. Statistical significance was assumed at p < 0.05.

qPCR of ChAT/vAChT mRNA against housekeeping in LA-N-2, p-values are found in Results (Welch two-sample t-test): Chat, Day 2, 10 ng/ml, t = −3.2436, df = 4.9872; 100 ng/ml, t = −2.349, df = 4.1296; Day 3, 10 ng/ml, t = −2.8481, df = 6.9658; 100 ng/ml, t = −6.3786, df = 3.3998; Day 4, 100 ng/ml, t = −9.0836, df = 6.9835. vAChT, Day 2, t = 5.9222, df = 5.3619; Day 4, t = 7.1016, df = 4.8784. Number of biological replicates: 4.

qPCR of ChAT mRNA against housekeeping in LA-N-5, p-values are found in Figure S2 legend (Welch two-sample t-test): Day 2, 10 ng/ml, t = −4.5204, df = 3.059; Day 4, 10 ng/ml, t = −4.7639, df = 5.0369; 100 ng/ml, t = −4.9161, df = 2.0262. Number of biological replicates: 4.

Mean absolute count-change LA-N-5 vs LA-N-2, text of Figure 3 (Welch two-sample t-test): t = 2.6183, df = 1108. Number of compared miRs: 490.

Target count of families, Figure 5A: enriched in both vs. enriched in one cell, t = 3.1831, df = 73; enriched in mir-10 vs. enriched in other sex-independent families, t = −3.28, df = 7.1879. Number of compared families: 17 (5 vs 12).

Validation of miR-125b-5p targeting of AChE, p-values are found in Results (one-way ANOVA): Figure 6B, n = 4, f-ratio value 9.19882, df between treatments 2, within treatments 9; Figure 6C, control n = 5, 125b n = 4, 132 n = 3, f-ratio value 11.59814, df between treatments 2, within treatments 8. N refers to biological replicates.

## Supplemental Text and Figures

**Fig. S1.**
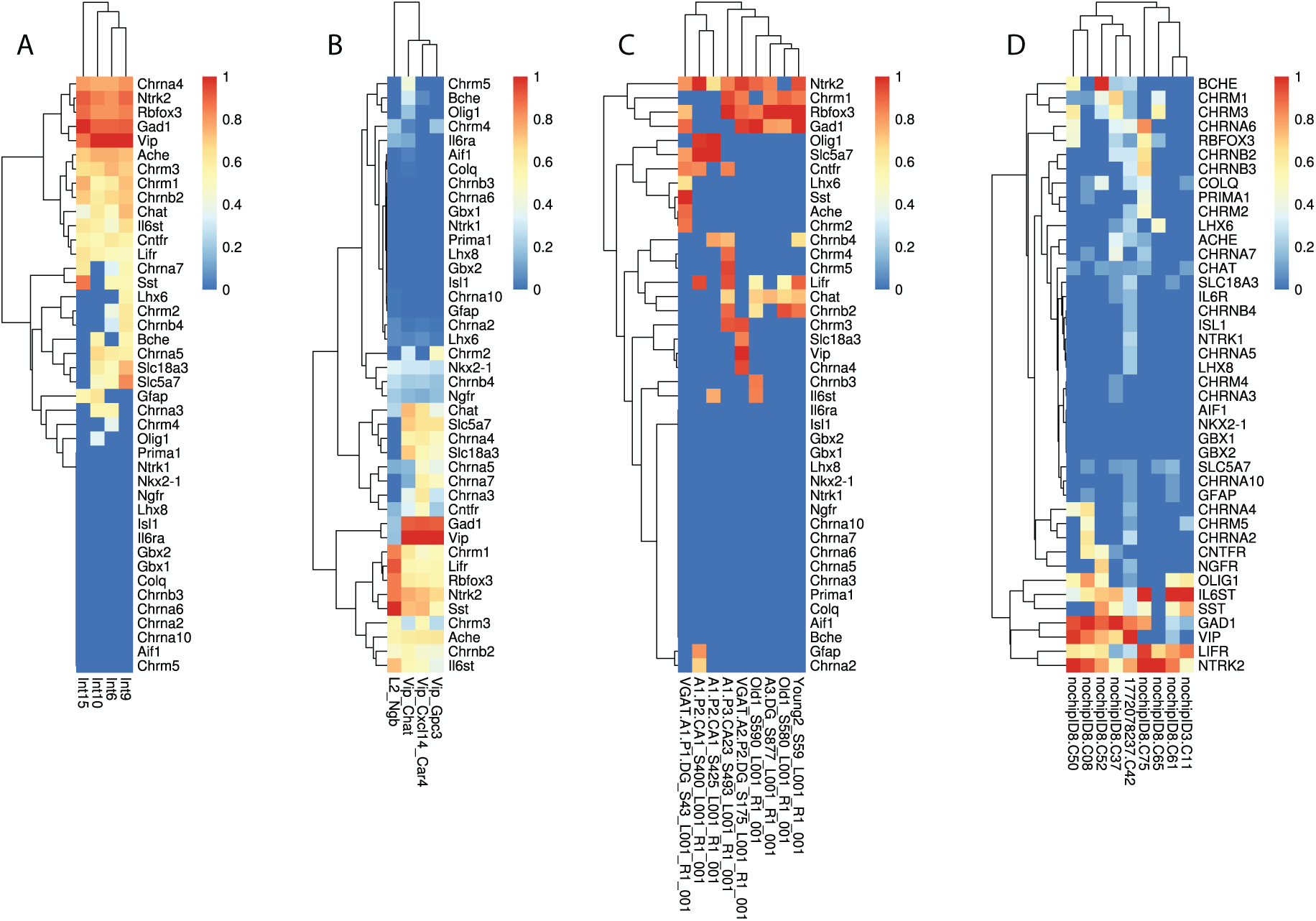
Single-cell sequencing expression heatmaps with original sample annotation from (A) Zeisel et al 2015, (B) Tasic et al 2016, (C) Habib et al 2016, (D) Darmanis et al 2015, related to Figure 3.

**Fig. S2.**
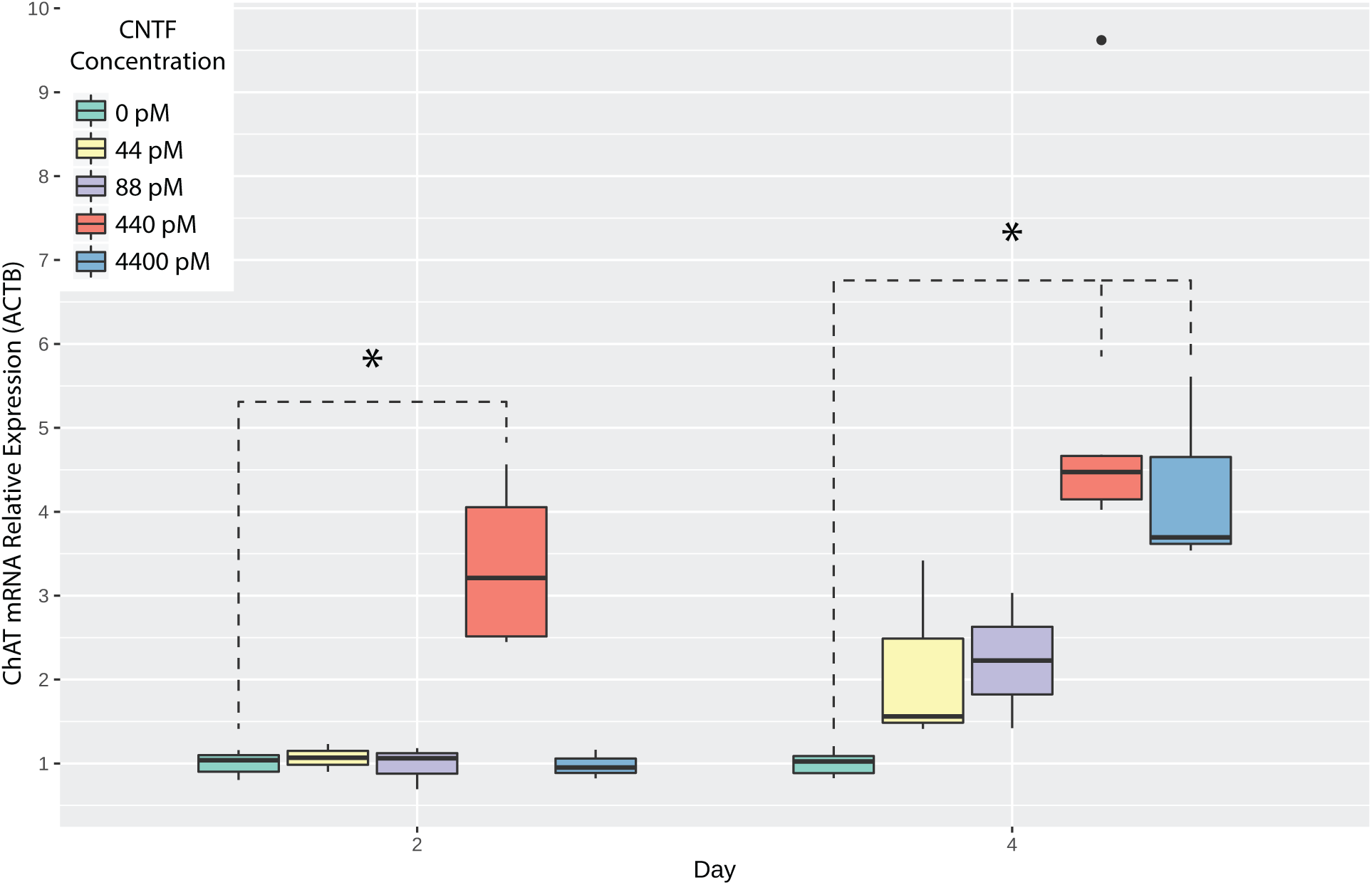
Dose-response-curve of LA-N-5 during CNTF-induced cholinergic differentiation as measured by expression of CHAT mRNA relative to ACTB. Significant differences at 10 ng/ml CNTF after 2 days (p = 0.019) and 4 days (p = 0.005), and 100 ng/ml after 4 days (p = 0.038). Related to Figure 4.

**Fig. S3.**
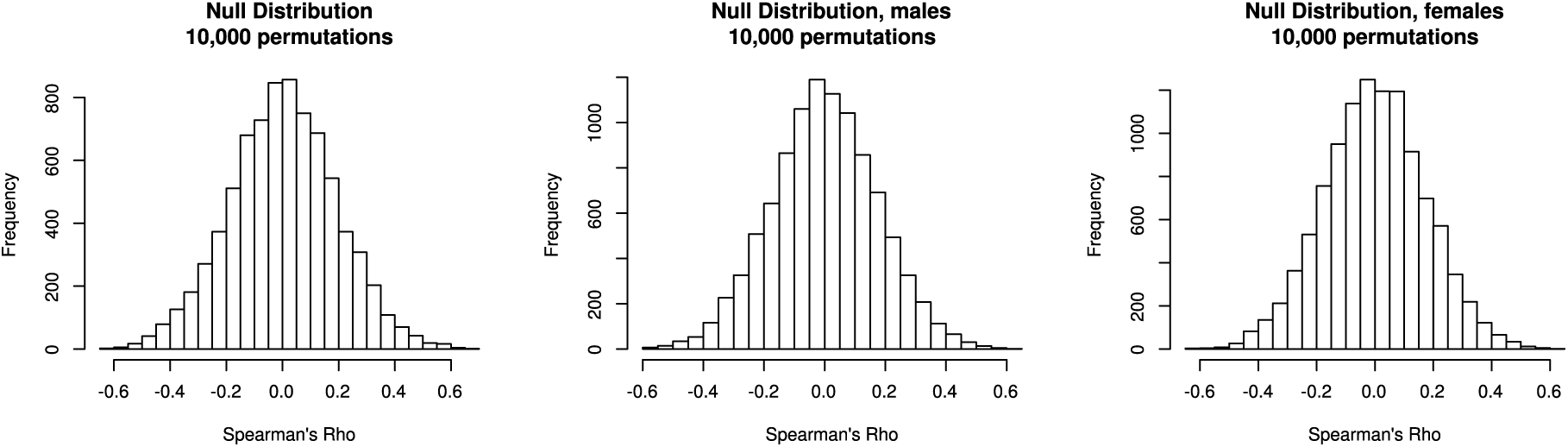
Unbiased meta-analysis null distributions of Spearman’s rho in sex-independent, male, and female datasets, related to STAR Methods - Whole transcriptome meta-analysis, Figure 1.

**Fig. S4.**
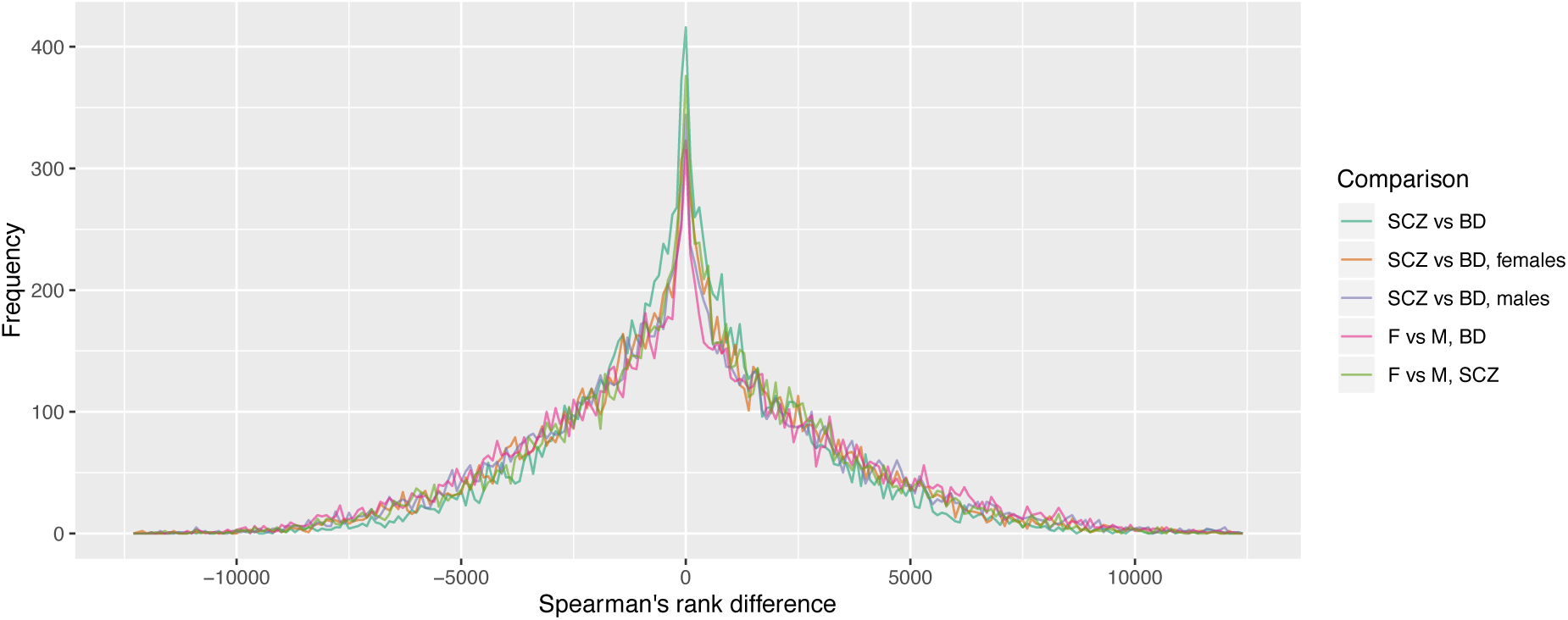
Spearman rank-differences between any two compared conditions in the GO-enrichment of beta-values from unbiased meta-analysis, related to STAR Methods - Whole transcriptome meta-analysis, Figure 1.

**Fig. S5.**
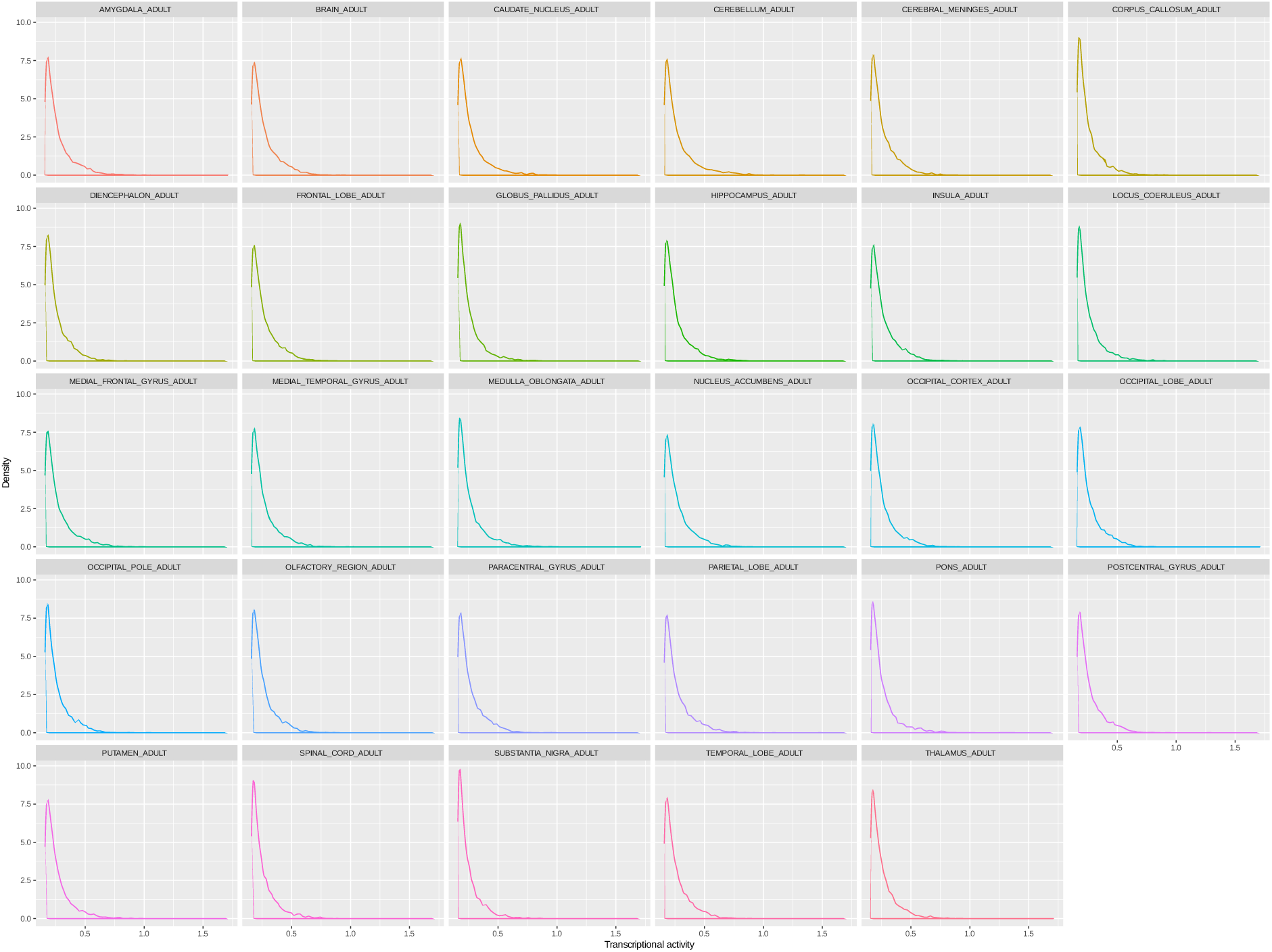
Density plots of transcriptional activities in analysed CNS brain regions derived from the dataset of Marbach et al 2016 (top 1% most active transcription factors in each brain region), related to STAR Methods - miR-gene-TF targeting.

